# Multilevel regulation of NF-κB signaling by NSD2 suppresses *Kras*-driven pancreatic tumorigenesis

**DOI:** 10.1101/2023.09.11.557111

**Authors:** Wenxin Feng, Ningning Niu, Ping Lu, Hanyu Rao, Zhuo Chen, Wei Zhang, Chunxiao Ma, Changwei Liu, Yue Xu, Wei-Qiang Gao, Jing Xue, Li Li

**Author notes:** **Corresponding authors Dr. Li Li** State Key Laboratory of Systems Medicine for Cancer, Renji-Med X Clinical Stem Cell Research Center, Ren Ji Hospital, School of Biomedical Engineering & Med-X Research Institute, Shanghai Jiao Tong University, 1954 Huashan Rd, Shanghai, China., **Dr. Jing Xue**, State Key Laboratory of Systems Medicine for Cancer, Stem Cell Research Center, Ren Ji Hospital, School of Medicine, Shanghai Jiao Tong University, 160 Pujian Rd, Shanghai 200127, China. equal contribution (Wenxin Feng, Ningning Niu).

## Abstract

Pancreatic ductal adenocarcinoma (PDAC) is a clinically challenging cancer with a dismal overall prognosis. NSD2 is an H3K36-specific di-methyltransferase which has been reported to play a crucial role in promoting tumorigenesis. Here, we demonstrate that NSD2 acts as a putative tumor suppressor in *Kras*-driven pancreatic tumorigenesis. Low level of NSD2 indicates aggressive feature of PDAC. NSD2 restrains the mice from inflammation and *Kras*-induced ductal metaplasia, while NSD2 loss facilitates pancreatic tumorigenesis. Mechanistically, NSD2-mediated H3K36me2 promotes the expression of IκBα, which inhibits the phosphorylation of p65 and NF-κB nuclear translocation. More importantly, NSD2 interacts with the DNA binding domain of p65, attenuating NF-κB transcriptional activity. Furthermore, inhibition of NF-κB signaling relieves the symptoms of *Nsd2*-deficient mice. Together, our study reveals the important tumor suppressor role of NSD2 and multiple mechanisms by which NSD2 suppresses both p65 phosphorylation and downstream transcriptional activity during pancreatic tumorigenesis. This study contributes to understanding the pathogenesis of pancreatic tumorigenesis and identifies a novel negative regulator of NF-κB signaling.

## Introduction

Pancreatic ductal adenocarcinoma (PDAC) is a deadly disease that is estimated to become the second-leading cause of cancer death in the US by 2030 at a rate of 0.5% to 1.0% per year ^1,2^. In 90% of cases, pancreatic cancer is discovered at a late stage, when it has spread beyond the pancreas, and over half have metastasized to other organs ^2,3^. Multiple genetic alterations are involved in the pathophysiology of PDAC, including *KRAS*, *TP53*, *SMAD4* and *CDKN2A* ^4^, and *KRAS* variants are identified in 90% to 92% of individuals with PDAC ^4^. Mouse models expressing *Kras* variants in acinar cells typically develop distinct precursor lesions known as pancreatic intraepithelial neoplasia (PanIN).

Nuclear factor kappa-light-chain-enhancer of activated B cells (NF-κB) is a transcription factor and key inducer of inflammatory responses ^5^. One of the major subunits of NF-κB, RelA (p65), forms heterodimers with the structurally related p50 protein ^6^. In the canonical pathway, most p65-p50 heterodimers (NF-κB) are located within the cytoplasm under basal conditions due to its association with inhibitors of NF-κB, IκBα. Stimulation of cells with NF-κB-activating ligands such as the cytokine tumor necrosis factor (TNF) results in the degradation of IκBα, and then NF-κB translocates to the nucleus, where it regulates transcriptional programs ^7^. Studies have shown that NF-κB signaling is constitutively activated in almost 70% of pancreatic cancer specimens ^8^ and plays a crucial role in PDAC development, progression and resistance ^9^.

Pancreatic tumorigenesis is also mediated by epigenetic regulation, which includes DNA methylation, histone modifications and chromatin remodeling ^10–12^. NSD2 is a histone methyltransferase that can catalyze the di-methylation of histone H3 at lysine 36 (H3K36me2), a permissive mark associated with active gene transcription ^13–15^. NSD2 has been reported to be amplified, mutated, or overexpressed in human cancers. The NSD2 t (4;14) chromosomal translocation in multiple myeloma (MM) is associated with the overexpression of NSD2 and leads to a poor prognosis ^16,17^. NSD2 is overexpressed in invasive prostate cancer, especially in metastases, and is associated with the poor prognosis of prostate cancer patients ^18^. Similar results have also been reported in other tumor types, such as esophageal carcinoma, stomach carcinoma, hepatocellular carcinoma, lung cancer, and corpus uteri malignancy, etc.^15^. Many studies have identified that NSD2 promotes cell proliferation, migration, invasion and epithelial-mesenchymal transformation (EMT), proving that it plays a crucial oncogenic role ^13–15^. However, the mechanism of NSD2 in pancreatic tumorigenesis remains unclear.

In this study, we investigated the role of NSD2 in *Kras*-induced pancreatic tumorigenesis. We found that NSD2 overexpression inhibited inflammation and *Kras*-induced ductal metaplasia in mice, whereas NSD2 loss facilitated *Kras*-induced tumorigenesis. Our results demonstrate that NSD2-mediated H3K36me2 promotes the expression of IκBα, inhibiting the phosphorylation of p65. More importantly, NSD2 interacts with the DNA binding domain of the NF-κB p65 protein, attenuating NF-κB transcriptional activity. Therefore, we uncovered multilevel regulation of NF-κB signaling by NSD2 during pancreatic tumorigenesis.

## Results

### The low protein level of NSD2 indicates aggressive feature of PDAC

Considering that NSD2 plays an oncogenic role in many tumor types, we first evaluated the association between *NSD2* expression and survival in pan-cancer based on the TCGA database. The results revealed that patients with elevated *NSD2* expression (top 50%, n=4751) had significantly shorter overall survival and disease-free survival than patients with low *NSD2* level (bottom 50%, n=4751). However, survival analysis of PDAC patients from the TCGA dataset indicated that *NSD2* mRNA level did not predict overall survival or disease-free survival.

Next, we examined the *NSD2* mRNA level in public human PDAC datasets and found that the expression level of *NSD2* in PDAC samples was lower in dataset GSE28735 ^19,20^, while it was higher in dataset GSE15471 ^21^. In addition, the *NSD2* expression level did not change in GSE16515 ^22^ (Fig. 1c).

**Fig 1.**
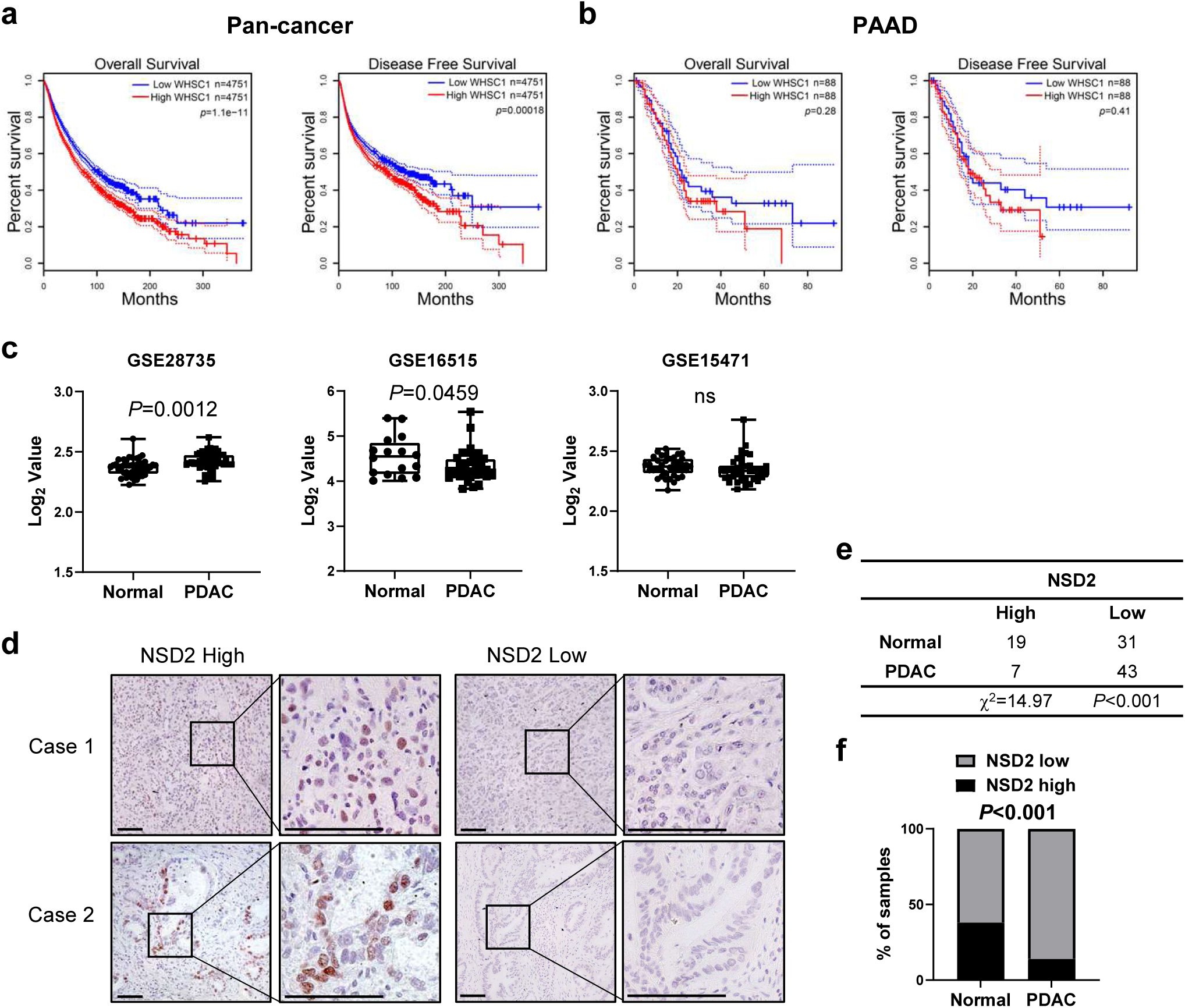
Low protein level of NSD2 indicates aggressive feature of PDAC. **a,** Kaplan-Meier overall survival (left) and disease-free survival (right) curves according to NSD2 expression were analyzed using TCGA pan-cancer database (high WHSC1 expression, 4751 cases, 50%; low WHSC1 expression, 4751 cases, 50%). **b,** Kaplan-Meier overall survival (left) and disease-free survival (right) curves according to NSD2 expression were analyzed using TCGA PAAD database (high WHSC1 expression, 88 cases, 50%; low WHSC1 expression, 88 cases, 50%). **c,** Box plot of NSD2 mRNA level in pancreas controls and PDAC specimens (using dataset GSE28735, GSE16515 and GSE15471). In boxplots (middle line depicts the median and the whiskers the min-to-max range). **d,** Representative IHC staining indicates high and low expression of NSD2 in PDAC tissue array. Scale bars: 100 μm. **e,** Statistics of high and low expression of NSD2 in pericarcinomatous samples and PDAC samples. **f,** NSD2 expression in pericarcinomatous samples and PDAC samples is quantified (χ2 test).

Given the ambiguous results of *NSD2* mRNA level in PDAC datasets, we further assess the clinical relevance of NSD2 protein level in a PDAC tissue array. We performed immunohistochemistry (IHC) analyses to determine the NSD2 level in a PDAC tissue array (Fig. 1d), and the quantification results showed that the NSD2 level in the PDAC samples was lower than that in the pericarcinomatous samples (Fig. 1e-f). However, there is no significant difference in H3K36me2 level between the PDAC samples and the pericarcinomatous samples in the tissue array cohorts (Fig. S1b-d). Taken together, the low NSD2 protein level, rather than H3K36me2 level, indicates aggressive feature of PDAC.

### NSD2 overexpression restrains pancreatic ductal metaplasia induced by inflammation and / or *Kras* mutation

To assess the role of NSD2 in pancreatic tumorigenesis, we generated pancreatic-specific NSD2-overexpressing mice together with oncogenic *Kras* mutation. Mice harboring a single copy of a minigene consisting of a CAGGS (a hybrid chicken β-actin and cytomegalovirus) promoter, a loxP-STOP-loxP (LSL) cassette, and Myc-tagged *Nsd2* cDNA knocked into the Rosa26 locus (*Nsd2*^OE/+^) ^18^ were crossed with *Pdx*^cre^; LSL-*Kras*^G12D^ mice (referred to as PK mice) to obtain *Pdx*^cre^; LSL-*Kras*^G12D^; *Nsd2*^OE/+^ mice (referred to as PKN^O^ mice) (Fig. 2a). The overexpression of NSD2 in the PKN^O^ pancreas was verified by using Western blotting and IHC (Fig. 2b-c). To investigate the biological function of NSD2 in the pancreas under the context of injury, we first applied a caerulein-induced pancreatitis recovery model (RAP) to 8-week-old PK and PKN^O^ mice (100 μg/kg, 8 hourly injection/day) (Fig. 2d). In comparison with PK mice, there were much less acinar-to-ductal metaplasia (ADM) lesions in the pancreata of PKN^O^ mice on both day 7 and day 18, demonstrating that NSD2 overexpression significantly restrained the inflammation-induced ADM (Fig. 2e-f). In addition, we compared the ductal lesions between PK and PKN^O^ mice at 24 weeks of age to illustrate the impact of NSD2 overexpression on *Kras*-induced ductal metaplasia. Strikingly, PKN^O^ mice showed significantly reduced development of ADM, PanIN and PDAC lesions compared to PK mice (Fig. 2g-h). Consistently, CK19^+^ and Ki67^+^ signals decreased dramatically in PKN^O^ lesions (Fig. 2i). Furthermore, pancreatic acinar explants derived from PKN^O^ mice dedifferentiated less into ductal-like structures than explants from PK mice in response to transforming growth factor α (TGFα) (Fig. 2j-k). Collectively, these findings indicate that NSD2 overexpression reinforces the acinar homeostasis and inhibits the pancreatic ductal metaplasia induced by both inflammation and *Kras*^G12D^.

**Fig 2.**
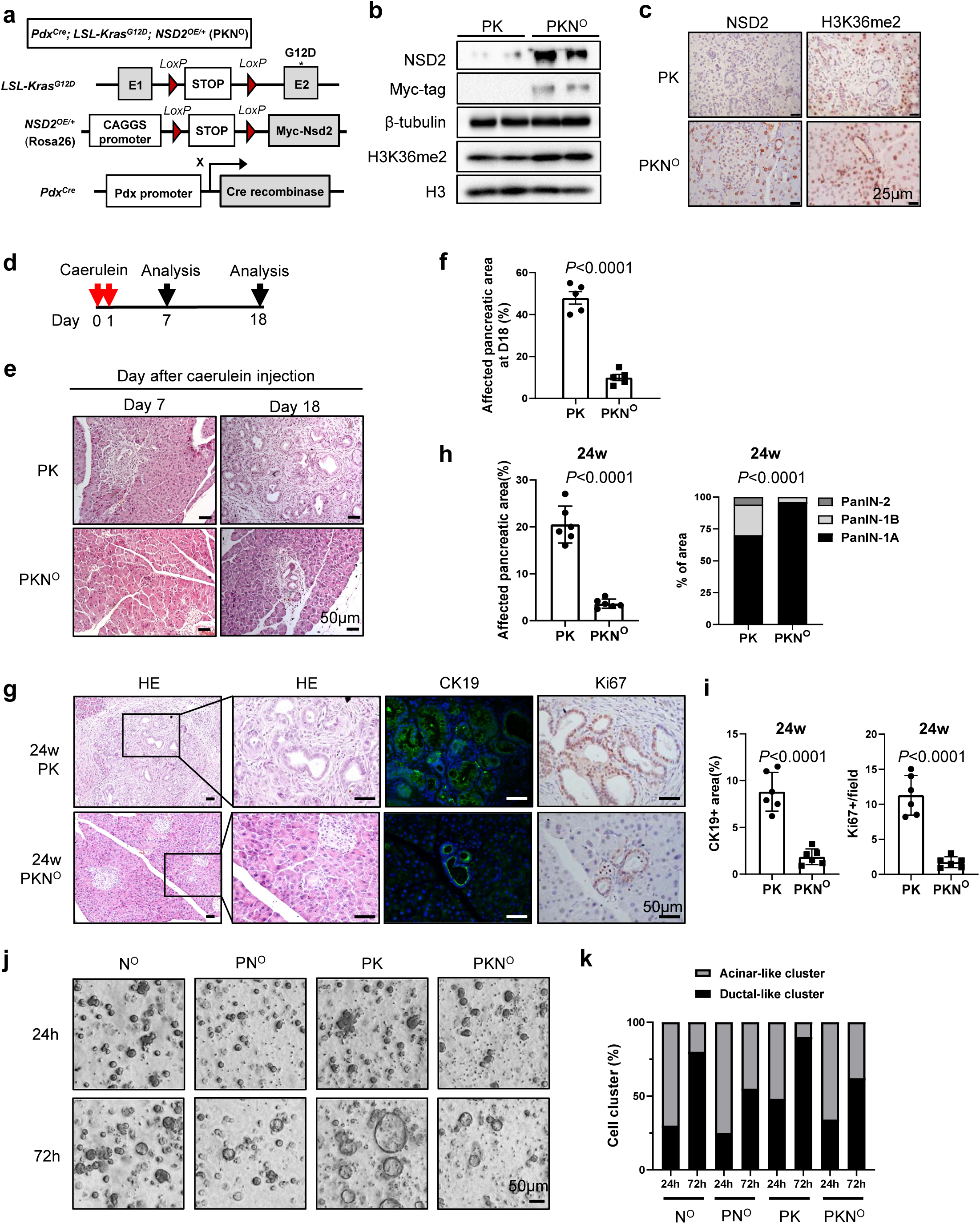
NSD2 overexpression restrains pancreatic ductal metaplasia induced by inflammation and / or *Kras* mutation. **a,** Construction and breeding strategy of *Pdx*^cre^; LSL-*Kras*^G12D^; *Nsd2*^OE/+^ mice (PKN^O^). **b,** Western blotting analysis of NSD2 and H3K36me2 expression in pancreas tissues of PK and PKN^O^ mice. Experiments were repeated at least three times, with similar results. **c,** IHC analysis of NSD2 and H3K36me2 expression in pancreas tissues of PK and PKN^O^ mice. Experiments were repeated at least three times, with similar results, and representative images are shown. Scale bars: 25 μm. **d,e,** PK and PKN^O^ mice were administrated with caerulein (100 µg/kg, 8 hourly injection/day) for consecutive 2 days. Pancreas tissues were collected at indicated days for HE staining (n = 5 per group). **f,** Quantification of affected pancreatic area (%) in pancreas tissue from indicated mice at day 18 (n = 5 per group). **g,** Pancreatic tissues from indicated mice for staining of HE, CK19 and Ki67 (n = 6 per group). Scale bars: 50 µm. **h,** Quantification of affected pancreatic area (%) and ratio of PanIN-1A, PanIN-1B and PanIN-2 in affected pancreatic areas from indicated mice at 24 weeks (n = 6 per group). **i,** Quantification of CK19 positive area and Ki67 positive cells per field from indicated mice at 24 weeks (n = 6 per group). **j,k,** Representative images and statistic of acinar-to-ductal metaplasia of acinar explants on transforming growth factor α (TGFα) stimulation from indicated mice. *Nsd2*^OE/+^ referred to as N^O^. *Pdx*^cre^; *Nsd2*^OE/+^ mice referred to as PN^O^. Experiments were repeated at least three times, with similar results, and representative images are shown.

### NSD2 loss facilitates *Kras*-induced ductal metaplasia

We next aimed to determine whether NSD2 loss could promote pancreatic tumorigenesis. Previously described *Nsd2*^f/f^ mice ^18^ were crossed with *Pdx*^cre^; LSL-*Kras*^G12D^ mice (PK mice) to ablate NSD2 in the pancreas (*Pdx*^cre^; *Kras*^G12D^; *Nsd2*^f/f^, referred to as PKN^f/f^ mice). By using Western blotting, IHC and Real-Time quantitative PCR (RT-qPCR), we found that the H3K36me2 level was unchanged in the pancreas of PKN^f/f^ mice compared to PK mice, although *Nsd2* was thoroughly ablated (Fig. 3a-c). This finding was inconsistent with previous studies, which manifested that *Nsd2* deletion reduced H3K36me2 levels in prostate cancer ^18,23^, breast cancer ^24^, osteosarcoma ^25^, lung adenocarcinoma ^26^, etc. The maintained H3K36me2 level could be elucidated by the fact that the expression levels of demethylases of the H3K36 site, including *Kdm2b*, *Kdm4a*, *Kdm4b* and *No66*, were also significantly decreased in the *Nsd2*-deficient pancreas, while *Nsd1* and *Nsd3* expression remained unchanged (Fig. 3c).

**Fig 3.**
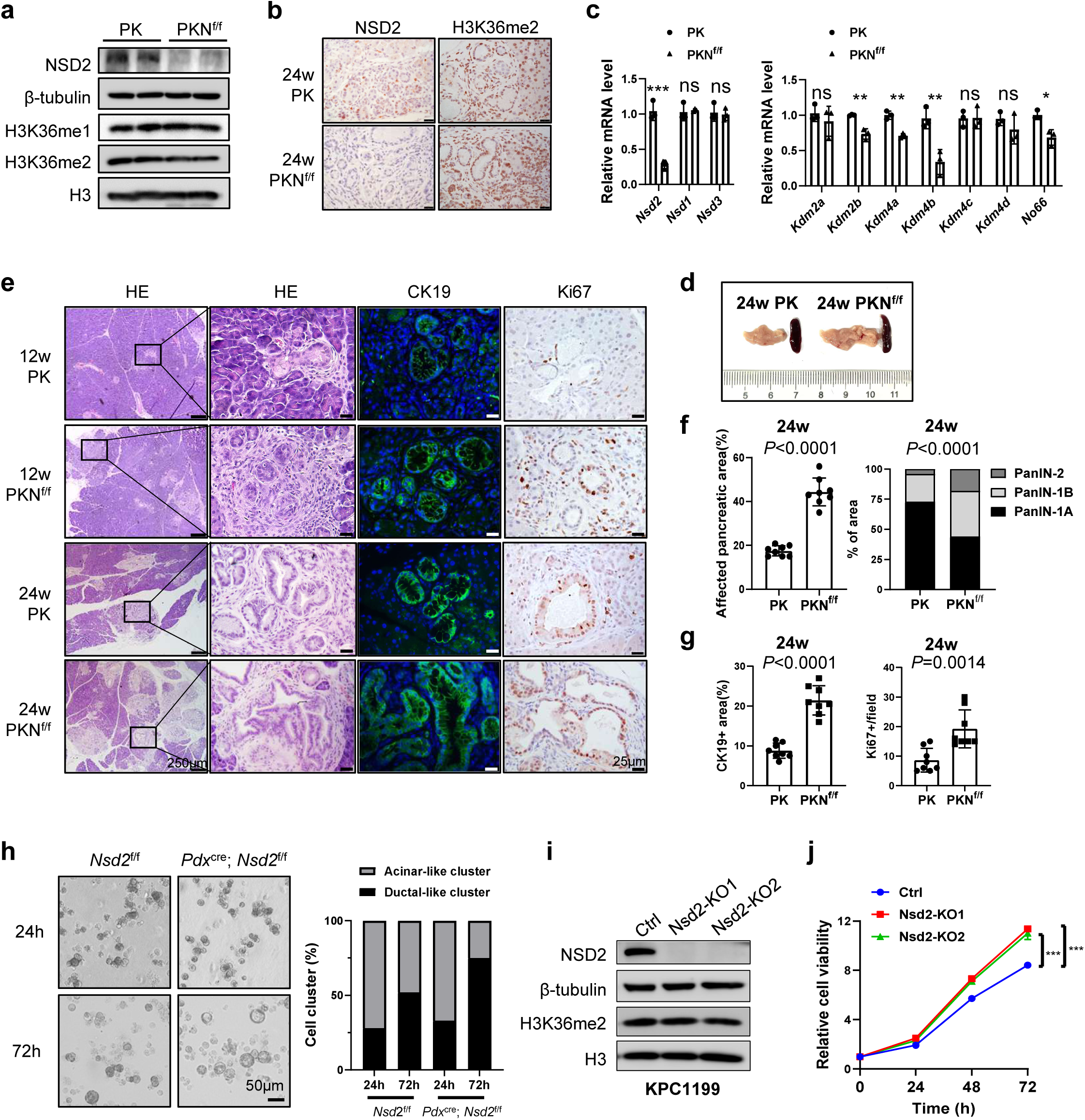
NSD2 loss facilitates *Kras*-induced ductal metaplasia. **a,** Western blotting analysis of NSD2, H3K36me1 and H3K36me2 expression in pancreas tissues of PK and PKN^f/f^ mice. Experiments were repeated at least three times, with similar results. **b,** IHC analysis of NSD2 and H3K36me2 expression in pancreas tissues of PK and PKN^f/f^ mice. Experiments were repeated at least three times, with similar results, and representative images are shown. Scale bars: 25 μm. **c,** RT-qPCR analysis of *Nsd2*, *Nsd1*, *Nsd3*, *Kdm2a*, *Kdm2b*, *Kdm4a*, *Kdm4b*, *Kdm4c*, *Kdm4d* and *No66* mRNA levels of PK and PKN^f/f^ mice. Experiments were repeated at least three times, with similar results. **d,** Representative macroscopic pancreas images of indicated mice. **e,** Pancreatic tissues from indicated mice for staining of HE, CK19 and Ki67 (n = 8 per group). Scale bars: 25 µm. **f,** Quantification of affected pancreatic area (%) and ratio of PanIN-1A, PanIN-1B and PanIN-2 in affected pancreatic areas from indicated mice at 24 weeks (n = 8 per group). **g,** Quantification of CK19 positive area and Ki67 positive cells per field from indicated mice at 24 weeks (n = 8 per group). **h,** Representative images and statistic of acinar-to-ductal metaplasia of acinar explants from indicated mice. Experiments were repeated at least three times, with similar results, and representative images are shown. **i,** Western blotting analysis of NSD2 and H3K36me2 expressions in Ctrl, Nsd2-KO1 and Nsd2-KO2 cells (KPC1199 derived). Experiments were repeated at least three times, with similar results. **j,** CCK8 assay of Ctrl, Nsd2-KO1 and Nsd2-KO2 cells (KPC1199 derived). Experiments were repeated at least three times, with similar results. PanIN, pancreatic intraepithelial neoplasia.

PK mice developed infrequent ADM lesions at 12 weeks of age and gradually progressed to low-grade panIN at approximately 24 weeks of age (Fig. 3d-e). In sharp contrast to age-matched PK mice, additional depletion of *Nsd2* accelerated the ductal metaplasia, and led to a more affected pancreatic area with much higher grades of PanIN/PDAC lesions at 24 weeks of age, along with enhanced cell proliferation (Fig. 3e-g). We also found that pancreatic acinar explants derived from *Pdx*^cre^; *Nsd2*^f/f^ mice dedifferentiated more into ductal-like structures than those from control mice (*Nsd2*^f/f^ mice) (Fig. 3h). Collectively, *Nsd2* loss synergized with oncogenic *Kras* to facilitate ADM transition and progression toward PanIN and PDAC.

To further analyze the effects of NSD2 loss, we depleted *Nsd2* with CRISPR/Cas9 in KPC1199, a murine pancreatic cancer cell line derived from KPC mice (*Pdx*^cre^; LSL-*Kras*^G12D^; LSL-*TP53*^R172H^). Western blotting analyses showed a reduction in NSD2 but not H3K36me2 (Fig. 3i). Consistent with the increased Ki67^+^ proportion, CCK8 assays confirmed the increased cell proliferation under NSD2 loss (Fig. 3j).

In line with the previous results (Fig 3c), the H3K36me2 level was also not impaired in *Nsd2*-deficient cells due to the decreased expression levels of the demethylases *Kdm2a*, *Kdm2b*, *Kdm4a*, *Kdm4b*, *Kdm4c* and *No66* (Fig. S2a). Together, these results demonstrate that *Nsd2* deletion promotes cell proliferation and accelerates pancreatic tumorigenesis.

### Loss of NSD2 activates the NF-κB signaling pathway

To gain mechanistic insight into how NSD2 loss promotes pancreatic tumorigenesis, we conducted an expression profile analysis of control (Ctrl) and Nsd2-KO cells by RNA sequencing (RNA-seq) (Fig. 4a). The standard for screening differentially expressed genes was FC>1.2 and P-value<0.05 was a significant differential gene. Gene Ontology (GO) analysis indicated that NSD2 loss significantly enriched the genes associated with the positive regulation of NF-κB transcription factor activities and activation of NF-κB-inducing kinase activities (Fig. 4b). To better understand NSD2-mediated signaling circuits, we performed GSEA, and the data showed that NSD2 loss significantly enriched the genes linked to EMT, TNFα signaling via NF-κB, E2F targets and inflammatory response (Fig. 4c).

**Fig 4.**
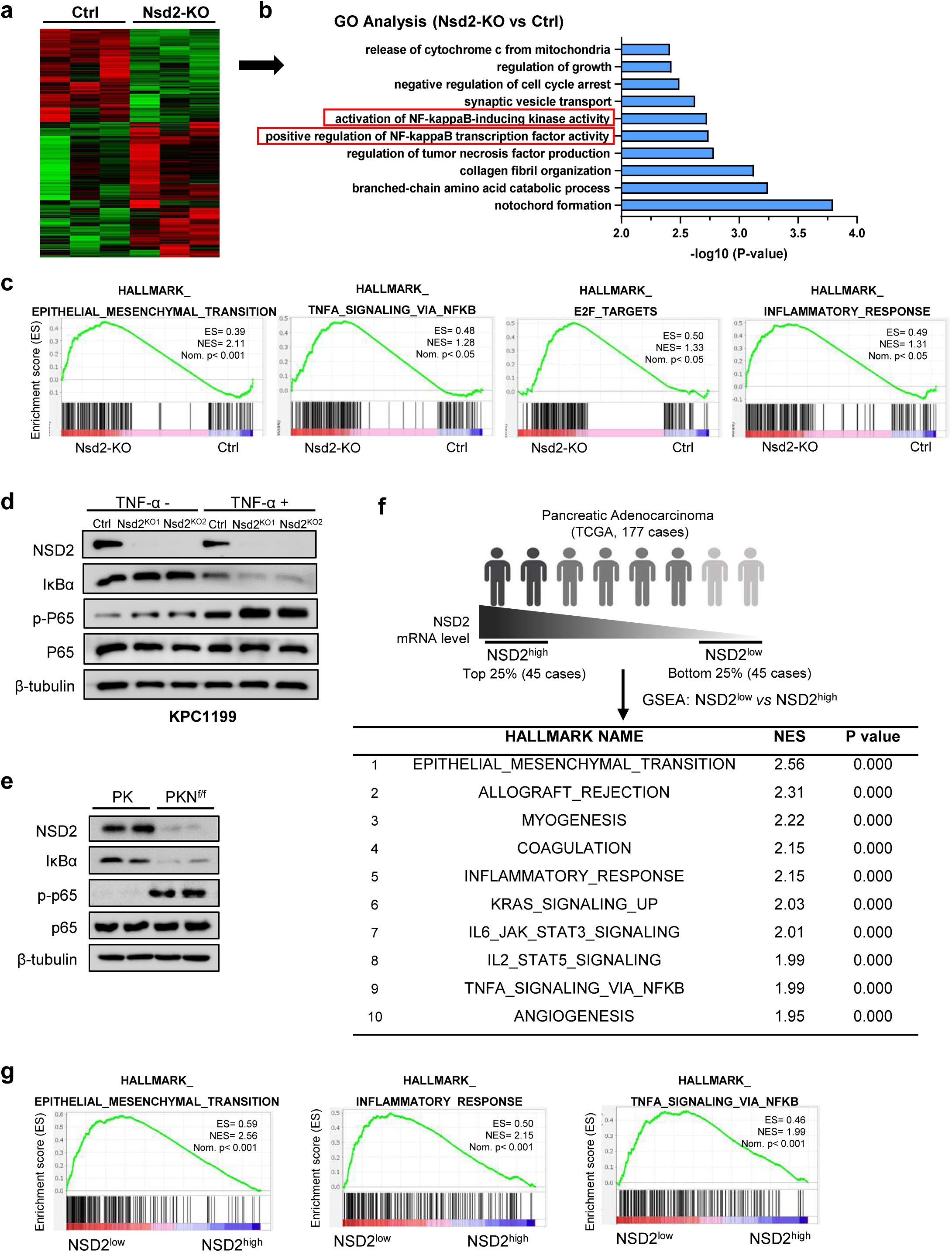
Loss of NSD2 activates NF-κB signaling pathway. **a,** Heat map of RNA-seq data to compare the gene expression of Ctrl and Nsd2-KO derived from KPC1199 cells. **b,** Go analysis of gene expression changes in RNA-Seq data (Ctrl and Nsd2-KO). **c,** GSEA enrichment plots of differentially expressed genes associated with *Nsd2* deletion. **d,** Western blotting analysis of NSD2, IκBα, p-p65 and p65 levels in Ctrl, Nsd2-KO1 and Nsd2-KO2 cells (KPC1199 derived) treated without or with TNF-α (20 ng/ml) for 30 min. Experiments were repeated at least three times, with similar results. **e,** Western blotting analysis of NSD2, IκBα, p-p65 and p65 expressions in empty vector (EV), Nsd2^WT^ and Nsd2^Y1179A^ cells (KPC1199 derived). Experiments were repeated at least three times, with similar results. **f,** GSEA analysis of gene expression between NSD2^low^ (patients with low expression of NSD2, 45 cases, 25%) and NSD2^high^ (patients with high expression of NSD2, 45 cases, 25%) from TCGA database. The significantly enriched hallmark pathways were listed. NES, normalized enrichment score. **g,** Epithelial Mesenchymal Transition, inflammatory response and TNF-α signaling via NF-κB enrichment plots in NSD2^low^ patients compared with NSD2^high^ patients are depicted.

To confirm that loss of NSD2 activates the NF-κB signaling pathway, Western blotting was performed to examine the protein levels of core members involved in canonical NF-κB signaling. The results showed that the phosphorylation of p65 was increased in Nsd2-KO cells, while the IκBα level decreased correspondingly (Fig. 4d). The Western blotting assays showed the same results in mice, which indicates the activation of the NF-κB signaling pathway in the pancreatic tissue of PKN^f/f^ mice (Fig. 4e). Together, these results prove that loss of NSD2 activates the NF-κB signaling pathway.

To assess the function of NSD2 in human PDAC, we compared the transcriptome differences between NSD2^high^ (top 25% of *NSD2* mRNA level, 45 cases) and NSD2^low^ (bottom 25% of *NSD2* mRNA level, 45 cases) patients in TCGA (PAAD) databases (Fig. 4f). We performed gene set enrichment analysis (GSEA) to gain a global view of the distinguished transcriptome profiles between NSD2^high^ and NSD2^low^ patients. Genes related to EMT, inflammatory response and TNFα signaling via NF-κB were significantly upregulated in PDAC with lower *NSD2* expression (Fig. 4f-g). Taken together, these findings revealed the link between NSD2 reduction and the activation of NF-κB signaling.

### NSD2-mediated H3K36me2 promotes *Nfkbia* expression

As shown in Fig. 4d-e, depletion of Nsd2 increased the phosphorylation of p65 and decreased the protein level of IκBα, suggesting that NSD2/H3K36me2 might positively regulate the expression of *Nfkbia*, IκBα coding gene. As expected, the *Nfkbia* mRNA level was downregulated in PKN^f/f^ mice, and upregulated in PKN^O^ mice (Fig. 5a). Downregulated *Nfkbia* expression was further validated in *Nsd2*-deficient KPC1199 cells (Fig. 5b). Furthermore, an NSD2 mutant vector for enzymatic activities (a tyrosine to alanine substitution at amino acid Y1179 abolished NSD2 methylation of nucleosomes) ^27^ was applied (Fig. 5c-d). The results showed that Nsd2^Y1179^ cells failed to upregulate *Nfkbia* expression level (Fig. 5e-f), indicating that the methyltransferase activity of NSD2 is essential to promote *Nfkbia* expression.

**Fig 5.**
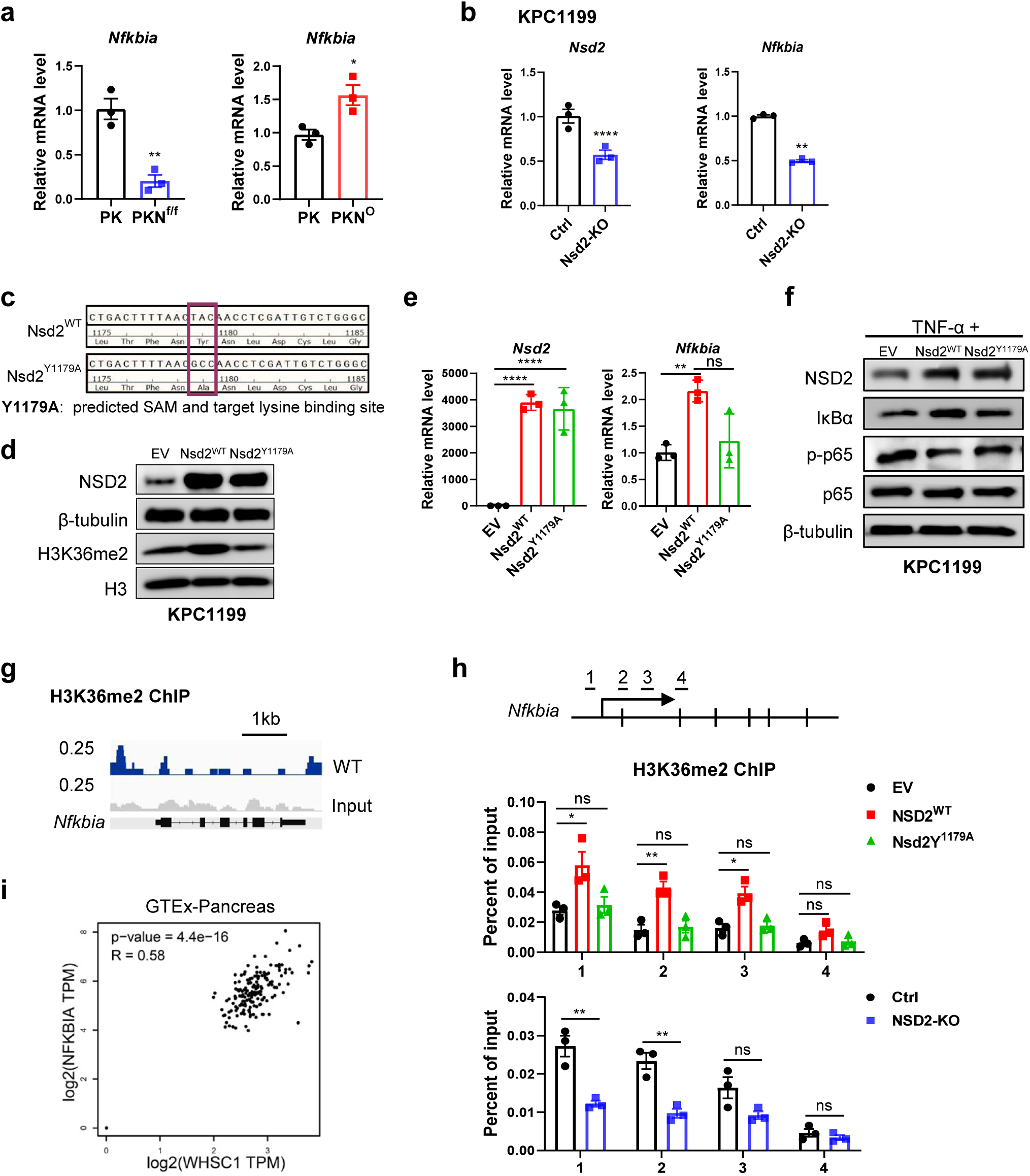
NSD2-mediated H3K36me2 promotes Nfkbia expression. **a,** RT-qPCR analysis of *Nfkbia* mRNA level of PK and PKN^f/f^ mice (left). RT-qPCR analysis of *Nfkbia* mRNA level of PK and PKN^O^ mice (right). Experiments were repeated at least three times, with similar results. **b,** RT-qPCR analysis of *Nsd2* and *Nfkbia* mRNA levels of Ctrl and Nsd2-KO cells (KPC1199 derived). Experiments were repeated at least three times, with similar results. **c,** Schematic diagram showing mutation site of NSD2. **d,** Western blotting analysis of NSD2 and H3K36me2 expressions in EV, Nsd2^WT^ and Nsd2^Y1179A^ cells (KPC1199 derived). Experiments were repeated at least three times, with similar results. **e,** RT-qPCR analysis of *Nsd2* and *Nfkbia* mRNA levels of EV, Nsd2^WT^ and Nsd2^Y1179A^ cells (KPC1199 derived). Experiments were repeated at least three times, with similar results. **f,** Western blotting analysis of NSD2, IκBα, p-p65, p65 levels in EV, Nsd2^WT^ and Nsd2^Y1179A^ cells (KPC1199 derived). Experiments were repeated at least three times, with similar results. **g,** Snapshot of H3K36me2 ChIP-Seq signals at the *Nfkbia* gene loci in KPC1199 cells. **h,** ChIP-qPCR of *Nfkbia* in EV, Nsd2^WT^ and Nsd2^Y1179A^ cells (KPC1199 derived) (middle). ChIP-qPCR of *Nfkbia* in Ctrl and Nsd2-KO cells (KPC1199 derived) (bottom). The location of the ChIP primer pairs used in the present study is denoted as numbers (top). Experiments were repeated at least three times, with similar results. **i,** The Pearson product-moment pair-wise gene correlation analysis between *NSD2* (*WHSC1*) and *NFKBIA* with GTEx-Pancreas expression database.

Although NSD2 loss did not significantly reduce the H3K36me2 level (Fig. 3a-b), we also examined the distribution patterns of H3K36me2. ChIP-seq was performed in KPC1199 cells, and the direct H3K36me2 occupancies within *Nfkbia* gene loci were observed, especially within the 5′-UTR and promoter regions (Fig. 5g). We further confirmed the existence of H3K36me2 marks at the promoter of *Nfkbia* gene by ChIP-qPCR, and the signals decreased along with *Nsd2* loss (Fig. 5h). Consistent with the previous results, the intensity of H3K36me2 bindings of the *Nfkibia* promoter increased in Nsd2^WT^ cells, rather than in Nsd2^Y1179^ cells (Fig. 5h). These data demonstrated that NSD2-mediated H3K36me2 directly regulates the expression of *Nfkbia.* Clinically, we found a strong correlation between *NSD2* and *NFKBIA* mRNA levels based on human GTEx-Pancreas database (Fig. 5i). Together, our results indicate that NSD2-mediated H3K36me2 promotes the expression of IκBα, which inhibits the phosphorylation of p65.

### *Nsd2* deletion promotes p65 nuclear translocation

Activation of the canonical NF-κB pathway leads to nuclear translocation of the p65-p50 dimer, which functions as a transcriptional activator ^5^. We next explored whether *Nsd2* deletion could promote p65 nuclear translocation. According to IHC staining, more p65 proteins were accumulated in the nucleus in the pancreatic tissue of PKN^f/f^ mice than in that of PK mice (Fig. 6a). Moreover, compared with the Ctrl, two independent Nsd2-KO cell lines both exhibited more nuclear translocation of p65 in response to TNF-α treatment (Fig. 6b). In addition, immunofluorescence (IF) analysis also revealed that loss of Nsd2 promoted nuclear translocation of p65 (Fig. 6c). These data all indicate that *Nsd2* deletion results in elevated p65 nuclear accumulation, which further demonstrates the enhanced activation of NF-κB signaling.

**Fig 6.**
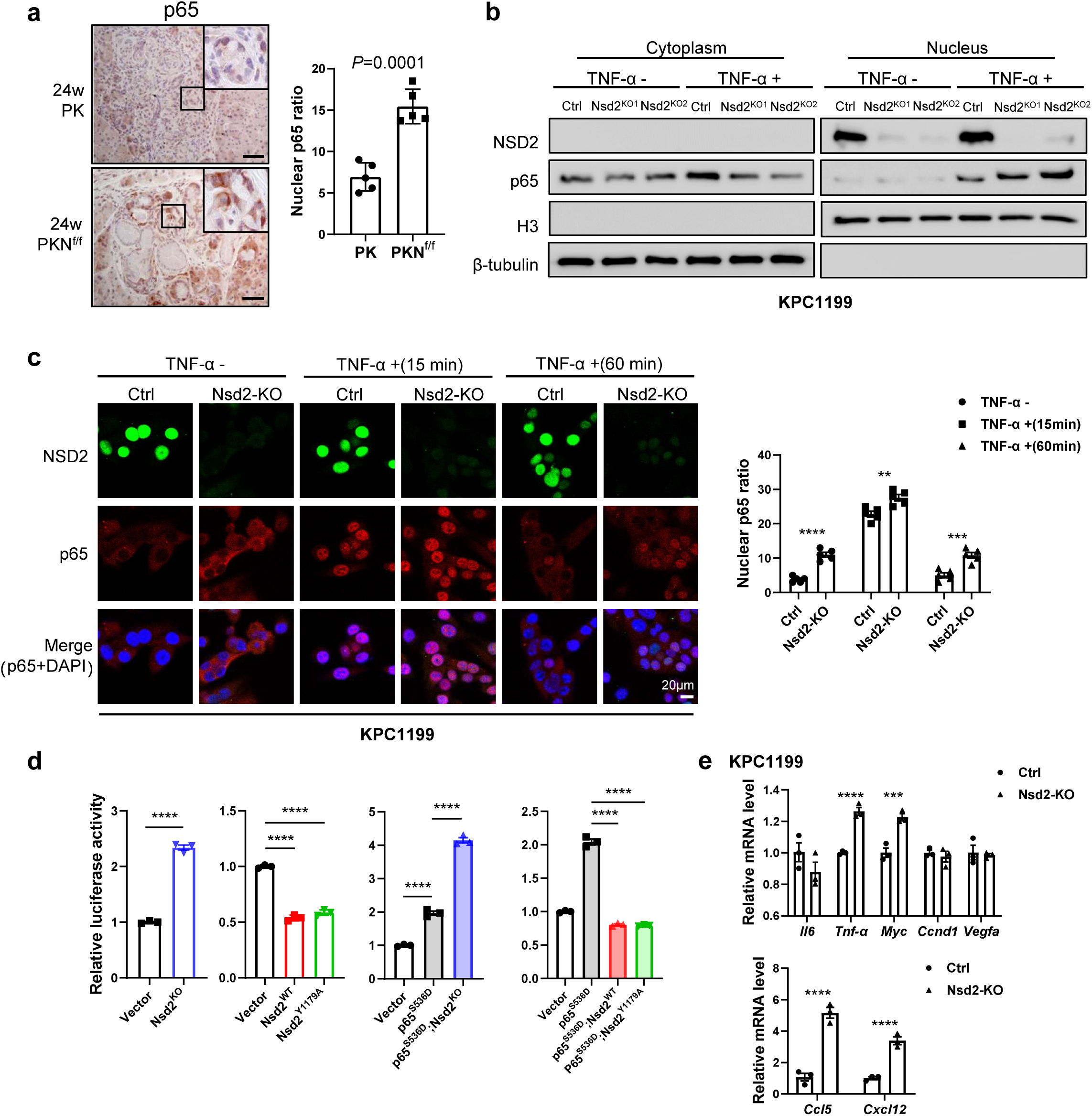
*Nsd2* deletion promotes p65 nuclear translocation. **a,** IHC analysis of p65 expression in pancreatic tissues from indicated mice at 24 weeks (n = 5 per group). Quantification of nuclear p65 mean intensity is shown on the right. Scale bars: 50 μm. **b,** Western blotting analysis of NSD2 and p65 in cytoplasmic and nuclear fractions in Ctrl, Nsd2-KO1 and Nsd2-KO2 cells (KPC1199 derived) treated without or with TNF-α (20 ng/ml) for 30 min. Experiments were repeated at least three times, with similar results. **c,** Immunofluorescence analysis of NSD2 and p65 expressions in Ctrl and Nsd2-KO cells (KPC1199 derived) treated without or with TNF-α (20 ng/ml) for 15 min and 60 min. Quantification of nuclear p65 mean intensity is shown on the right. Experiments were repeated at least three times, with similar results, and representative images are shown. **d,** Luciferase assays of NF-κB activity in 293T cells transfected with indicated plasmids. Experiments were repeated at least three times, with similar results. **e** RT-qPCR analysis of *Il6*, *Tnf-*α, *Myc*, *Ccnd1*, *Vegfa*, *Ccl5* and *Cxcl12* mRNA levels of Ctrl and Nsd2-KO cells (KPC1199 derived). Experiments were repeated at least three times, with similar results.

In addition, NF-κB luciferase reporter assays were performed in 293T cells. The plasmid expressing a dominant active form of p65 (p65^S536D^) was used to mimic phosphorylation at Ser536 of p65 ^28^. The results showed that NF-κB luciferase activity was significantly increased in the *Nsd2*-deficient cells but decreased in the *Nsd2*-overexpressing cells (both Nsd2^WT^ and Nsd2^Y1179^ cells) (Fig. 6d). RT-QPCR further validated that the expression levels of NF-κB target genes were significantly increased in Nsd2-KO cells, including *Tnf-*α*, Myc, Ccl5* and *Cxcl12* (Fig. 6e). These data demonstrate that loss of NSD2 increases NF-κB transcriptional activity, and NSD2 overexpression inhibits the activity. Notably, this effect is independent of NSD2 enzymatic activities.

### NSD2 interacts with the DNA binding domain of p65

As shown in Fig. 3a-b, H3K36me2 was not significantly reduced in *Nsd2*-deficient mice. In addition, luciferase assays also revealed that the ability of NSD2 to inhibit NF-κB transcriptional activity was independent of its enzymatic activities (Fig. 6d). Therefore, we next investigated whether NSD2-mediated NF-κB activation resulted from the physical interaction of NSD2 with p65. The localization of NSD2 and p65 was examined by IF staining in KPC1199 and PANC1 cells treated with or without TNF-α. The data showed that p65 was translocated to the nuclei 15 min after TNF-α treatment, and the co-localization between NSD2 and p65 occurred in the nucleus (Fig. 7a). We next assessed the endogenous interaction between NSD2 and p65 by co-immunoprecipitation (co-IP) with an anti-NSD2 antibody in KPC1199 and PANC1 cells treated with TNF-α. The results indicated that endogenous NSD2 showed a strong interaction with endogenous p65 (Fig. 7b). Exogenous co-IP assays also showed that both NSD2 and mutant NSD2^Y1179A^ were able to bind with P65 (Fig. 7c), suggesting that the protein interaction between NSD2 and p65 was independent of NSD2 enzymatic activities.

**Fig 7.**
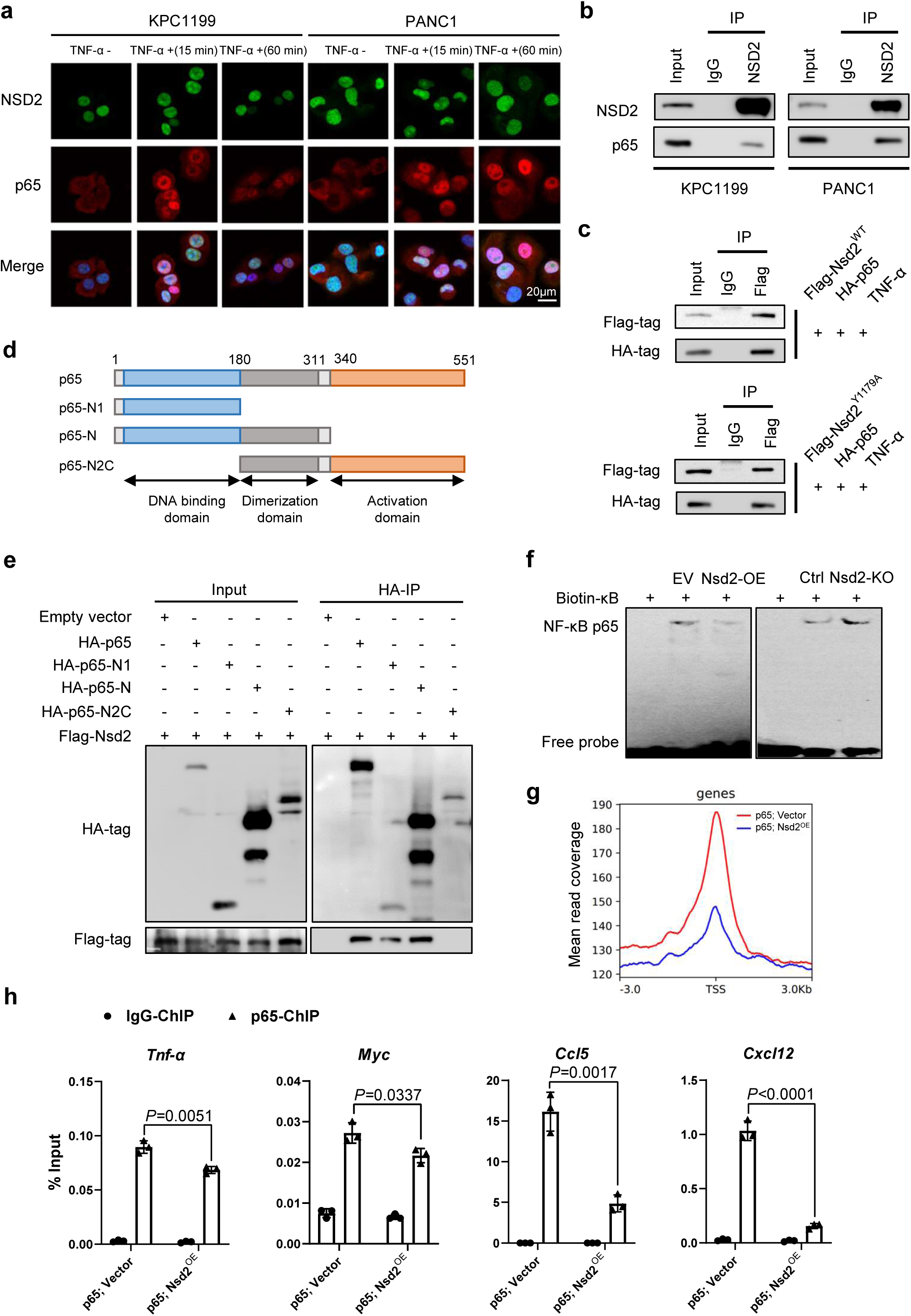
NSD2 interacts with the DNA binding domain of p65. **a,** Localization of NSD2 and p65 was observed by a confocal microscope. KPC1199 and PANC1 cells were treated without or with TNF-α (20 ng/ml) for 15 min and 60 min. Scale bars: 20 μm. Experiments were repeated at least three times, with similar results, and representative images are shown. **b,** NSD2 was immunoprecipitated from cells extracts of KPC1199 and PANC1 cells (treated with 20 ng/ml of TNF-α for 30 min) and immunoblotted with antibodies against NSD2 and p65. Experiments were repeated at least three times, with similar results. **c,** HEK293T cells were co-transfected with Flag-NSD2 and HA-p65, and then treated with 20 ng/ml of TNF-α for 30 min. Flag was immunoprecipitated from cells extracts and immunoblotted with antibodies against Flag and HA. Experiments were repeated at least three times, with similar results. **d,** Diagram of the truncation mutants of p65. The N-terminal DNA binding domain, dimerization domain and the activation domain are shown by blue, gray and yellow boxes, respectively. **e,** Flag-tagged NSD2 protein and HA-tagged wild-type and truncated p65 proteins were transiently expressed in HEK293T cells and and then treated with 20 ng/ml of TNF-α for 30 min. HA was immunoprecipitated from cells extracts and immunoblotted with antibodies against HA and Flag. Experiments were repeated at least three times, with similar results. **f,** EMSA assay for NF-κB p65 activity in KPC1199 cells transfected with indicated plasmids. Experiments were repeated at least three times, with similar results. **g,** Normalized read density of p65 ChIP-seq signals in KPC1199 cells transfected with indicated plasmids. **h,** ChIP-qPCR analysis of p65 binding for *Tnf-*α, *Myc*, *Ccl5* and *Cxcl12* promoter in KPC1199 cells transfected with indicated plasmids, and IgG was used as the control. Experiments were repeated at least three times, with similar results.

Next, we examined the NSD2-binding domain of p65 by co-IP with truncation mutants of p65 (Fig. 7d). Consistent with the data shown in Fig. 7c, NSD2 bound to full-length p65. Furthermore, the N-terminal domain contains the DNA binding domain of p65 (p65-N1 and p65-N), but not the dimer formation domain with the activation domain (p65-N2C), coprecipitated with flag-tagged NSD2, indicating that p65 interacts with NSD2 via its DNA binding domain (Fig. 7e). Therefore, we determined whether NSD2 affected the binding of NF-κB to the κB site *in vitro* by electrophoretic mobility shift assay (EMSA). NSD2 depletion enhanced NF-κB binding to the κB site, while NSD2 overexpression decreased it (Fig. 7f). Furthermore, we questioned whether NSD2 affected the binding of p65 to target gene promoters *in vivo* by chromatin immunoprecipitation (ChIP) followed by ChIP-seq assays. In line with our hypothesis, p65 peaks were enriched around the gene promoter region, and NSD2 overexpression resulted in a reduction in the intensity of p65 signals (Fig. 7g). ChIP–qPCR assays were used to validate the p65 binding at the promoters of the *Tnf-*α*, Myc, Ccl5 and Cxcl12* genes, and the results showed that the binding of p65 to these genes was significantly decreased along with the NSD2 overexpression (Fig. 7h). Collectively, these results indicate that NSD2 interacts with the DNA binding domain of p65 and suppresses the binding of NF-κB to the target genes.

### Inhibition of NF-κB signaling relieved the symptoms of *Nsd2*-deficient mice

Given that loss of NSD2 facilitates *Kras*-induced ductal metaplasia by activating the NF-κB signaling pathway, we investigated whether NF-κB signaling inhibitors could rescue the mice from these symptoms. JSH-23 and SN-50, specific NF-κB inhibitors that directly inhibit NF-κB nuclear transport ^29–34^ were utilized in 12-week-old PK and PKN^f/f^ mice. The compound was administered by intraperitoneal injection (i.p.) four times a week for four weeks, respectively (Fig. 8a). Compared with the vehicle control group, PKN^f/f^ mice treated with JSH-23 or SN-50 showed much fewer ADM/PanIN lesions (Fig. 8b-c), while PK mice treated with JSH-23 or SN-50 only showed a relatively modest decrease in ADM/PanIN lesions area (data not shown). In addition, the pancreas of treated mice exhibited significantly reduced CK19 and Ki67 (Fig. 8b). As expected, both JSH-23- and SN-50-treated mice displayed decreased nuclear p65 staining signals (Fig. 8b) and p65 target genes expression levels (Fig. 8d). Hence, inhibition of NF-κB signaling relieved the symptoms caused by *Nsd2* deletion in mice. We also treated the *Nsd2*-deficient KPC1199 cells with JSH-23 and SN-50 respectively. As shown by CCK8 assays, *Nsd2* loss promoted cell proliferation, which was significantly inhibited by treatment with either JSH-23 or SN50 (Fig. 8e). Next, wound healing and transwell-based invasion assays showed decreased cell migration and invasion after treatment with either JSH-23 or SN50 (Fig. 8f-g). Collectively, these results suggested that pancreatic tumorigenesis *in vivo* and cell proliferation *in vitro* due to *Nsd2* deletion are mediated by the activation of NF-κB signaling.

**Fig 8.**
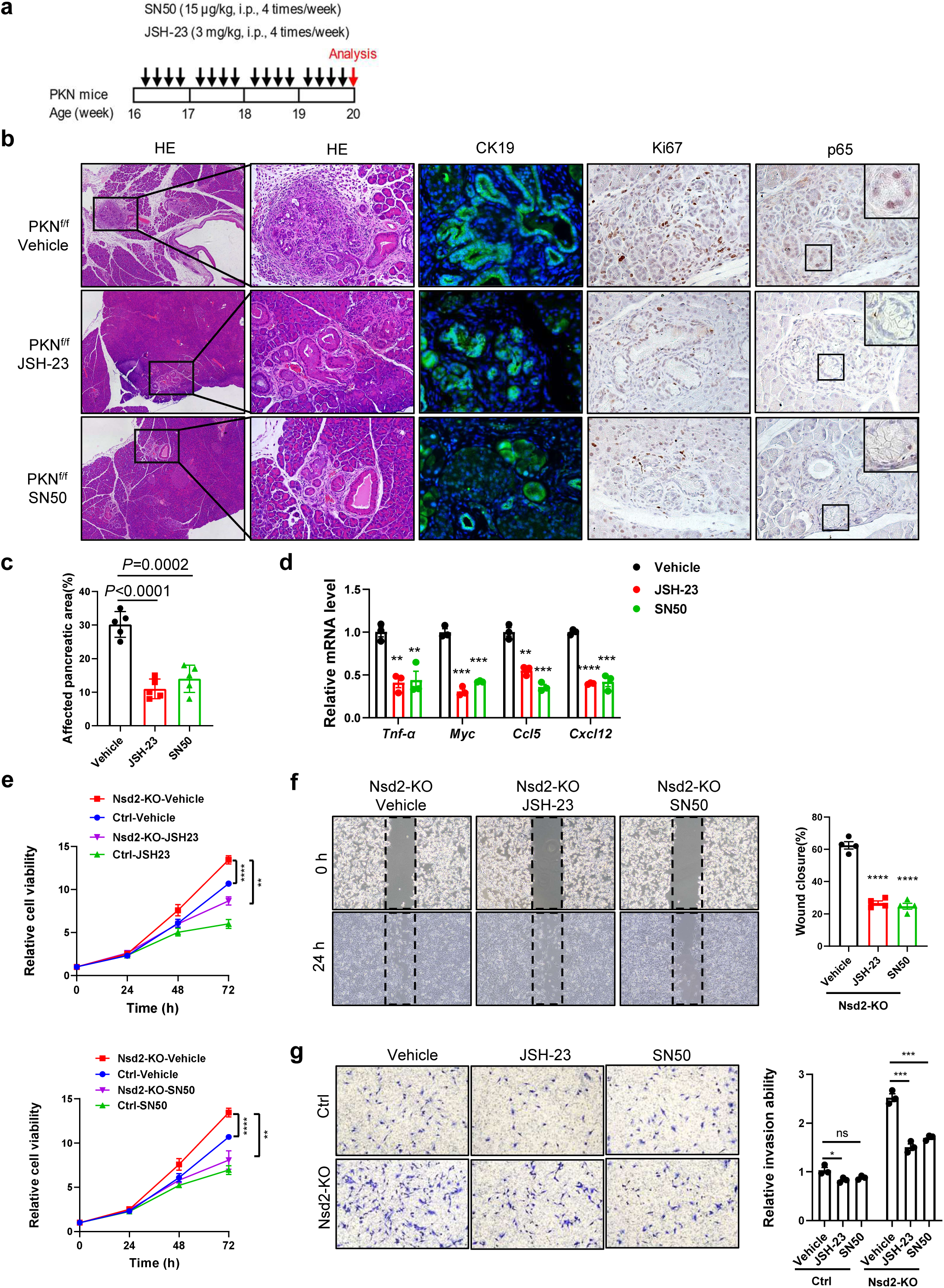
Inhibition of NF-κB signaling relieves the symptom of *Nsd2*-deficient mice. **a,** Scheme of treatment given for each injection. **b,** Pancreatic tissues from PKN^f/f^ mice treated with JSH-23, SN50 or vehicle control for staining of HE, CK19, Ki67 and p65 (n = 5 per group). **c,** Quantification of the affected pancreatic area in PKN^f/f^ mice treated with JSH-23, SN50 or vehicle control (n = 5 per group). **d,** RT-qPCR analysis of *Tnf-*α, *Myc*, *Ccl5* and *Cxcl12* mRNA levels of PKN^f/f^ mice treated with JSH-23, SN50 or vehicle control. Experiments were repeated at least three times, with similar results. **e,** CCK8 assay of ctrl and Nsd2-KO KPC1199 cells treated with the NF-κB signaling inhibitor JSH-23, SN50 or vehicle control. Experiments were repeated at least three times, with similar results. **f,** Wound healing assay of Nsd2-KO KPC1199 cells treated with JSH-23, SN50 or vehicle control for 24 hours. Experiments were repeated at least three times, with similar results, and representative images are shown. **g,** Transwell-based invasion assay of ctrl and Nsd2-KO KPC1199 cells treated with JSH-23, SN50 or vehicle control for 24 hours. Experiments were repeated at least three times, with similar results, and representative images are shown.

## Discussion

PDAC is a clinically challenging cancer, due to both its late stage at diagnosis and its complex molecular pathology ^4^. Therefore, a new understanding of the pathogenesis of PDAC can help reveal new opportunities for early detection and effective targeted therapy of PDAC. NSD2 is an important histone methyltransferase ^13^, and its oncogenic role in many cancer types has been gradually revealed in recent years ^13–15^. In this study, we demonstrate that NSD2 overexpression restrains pancreatic tumorigenesis, while NSD2 loss facilitates it, by using genetically engineered mouse models (GEMMs) (Fig. 2-3). In line with *in vivo* data, mouse pancreatic acinar explants assays showed the same results (Fig. 2-3). Our results showed that NSD2 acted as a novel tumor suppressor in pancreatic cancer and revealed the multiple mechanisms by which NSD2 suppresses both p65 phosphorylation and downstream transcriptional activity during pancreatic tumorigenesis.

Recent work reported that NSD2-mediated H3K36me2 promoted EMT, tumor differentiation and metastasis in PDAC using cell lines and orthotopic implantation model ^35^. However, in contrast to cancer cell inoculation models, GEMMs develop *de novo* tumors in a natural immune-proficient microenvironment, closely mimicking the histopathological and molecular features of human PDAC. Our results showed that H3K36me2 was not significantly reduced in *Nsd2*-deficient mice, which could be due to the reduced expression of demethylases *Kdm2b, Kdm4a, Kdm4b* and *No66* (Fig. 3a-c). Moreover, NSD2 is a large protein of more than 150 kDa. Apart from the evolutionarily conserved catalytic SET domain, it contains other physiologically functional domains ^36^, which may play important roles in protein-protein and/or protein-DNA/RNA interaction ^37^. As expected, luciferase reporter assays and exogenous Co-IP assays revealed that NSD2 interacted with p65 and inhibited NF-κB transcriptional activity independently of its enzymatic activities (using a catalytic inactivating mutation NSD2^Y1179^ variant). As the function of NSD2 is largely dependent on its enzymatic activities, our results demonstrate that NSD2 inhibited NF-κB transcriptional activity in the nucleus by interacting with p65, which is a histone methylation-independent regulatory pathway.

During canonical NF-κB signaling, inflammatory stimulation activates the IKK complex, which phosphorylates the IκB, resulting in ubiquitination and proteasomal degradation of IκB ^38,39^. The degradation of IκB releases the p65/p50 NF-κB heterodimer, allowing its nuclear translocation and promoter binding for target gene transcription ^38,39^. Strict regulation of the NF-κB signaling pathway is critical for pathological inflammation and cancer development ^40^. It is necessary for negative regulatory mechanisms that attenuate extensive signaling activity for the initiation and propagation of NF-κB signaling. Studies of such negative regulation of NF-κB signaling have mostly focused on the reversal of ubiquitination in the cytoplasm mediated by deubiquitinating enzymes such as CYLD, OTULIN and A20 ^41–46^. A20 and CYLD inhibit NF-kB signaling by targeting TRAF6 upstream of IKK ^43,45^. However, there are only a few reports describing negative regulators of NF-κB signaling in the nucleus. For instance, PIAS1 ^47^ and Twist ^48^ can interact with p65 and repress the transcriptional activity of NF-κB. In this study, we reveal that NSD2 is another novel and physiologically important negative regulator of NF-κB in the nucleus and may act as a brake to control the NF-κB signaling.

In addition to being a histone methyltransferase, NSD2 can also methylate non-histone proteins as well. A previous study revealed that NSD2 directly methylated PTEN at K349 and enhanced the DNA damage repair ability in colorectal cancer ^49^. NSD2 was found to methylate Aurora kinase A (AURKA) at K14 and K117 and mediate cell proliferation via the p53 signal pathway ^50^. Another study also revealed that NSD2 methylated STAT3, which promoted the activation of the STAT3 pathway and enhanced the ability of tumor angiogenesis ^51^. In this study, we performed mass spectrometry (MS) analysis and found no evidence that NSD2 directly methylated p65 (data not shown). Our results demonstrated that NSD2 regulated p65 mainly through its protein interaction with p65 and occupying the DNA-binding domain of P65. However, whether NSD2 methylates or binds to other members of NF-κB needs to be investigated in the future.

In summary, we reveal the physiological significance of NSD2 in pancreatic tumorigenesis. NSD2 acts as a putative tumor suppressor in *Kras*-driven pancreatic tumorigenesis. NSD2-mediated H3K36me2 promotes the expression of IκBα, which inhibits the phosphorylation of p65 and NF-κB nuclear translocation. In the nucleus, NSD2 interacts with the DNA binding domain of p65, attenuating p65-mediated NF-κB transcriptional activity. Clinical data also support that the expression level of NSD2 is downregulated (Fig. 1) and negatively correlated with the NF-κB target genes (Fig. S4c) in human PDAC. Of note, the genes related to TNFα signaling via NF-κB were significantly enriched in the PDAC patients with lower NSD2 expression (Fig. 4f-g). This study reveals the multilevel regulation of NF-κB activity by NSD2 during pancreatic tumorigenesis. We have identified NSD2 as a key tumor suppressor during pancreatic tumorigenesis and as a novel negative regulator of NF-κB signaling. This study advances the understanding of the pathogenesis of pancreatic tumorigenesis and opens therapeutic opportunities for PDAC patients with *NSD2* loss/mutations by targeting the NF-κB signaling pathway.

## Methods

### Mice

NSD2 Flox mice (Nsd2^f/f^) and NSD2-overexpressing mice (Nsd2^OE/+^) were generated as previously reported^18^ and gifted by Prof. Jun Qin. *Pdx*^cre^ and LSL-*Kras*^G12D^ mice were purchased from The Jackson Laboratory. These strains were interbred to generate the experimental cohorts, which include the following genotypes: *Pdx*^cre^; *Nsd2*^f/f^ (PN^f/f^), *Pdx*^cre^; *Nsd2*^OE/+^ (PN^O^), *Pdx*^cre^; LSL-*Kras*^G12D^ (PK), *Pdx*^cre^; LSL-*Kras*^G12D^; *Nsd2*^f/f^ (PKN^f/f^), *Pdx*^cre^; LSL-*Kras*^G12D^; *Nsd2*^OE/+^ (PKN^O^). Mice were harvested at the indicated time for pancreas histology investigation. The mice that lose weight by more than 20% within 1 week will be euthanized and counted as death. All mice were bred and maintained at an animal facility under specific pathogen-free conditions. Mouse experimental protocols were approved by the Renji Hospital Animal Care and Use Committee (202201027).

### Human subjects

The human pancreatic tumor microarray from Ren Ji Hospital was approved with Local Ethics Committee approval and patient consent (KY2022-036-B). Briefly, clinical parameters of patients with pancreatic cancer were collected, including age, sex, stage, pathological diagnosis, differentiation status, TNM status and survival. None of the patients underwent preoperative chemotherapy or radiation therapy prior to surgery.

### Experimental pancreatitis recovery model

Experimental pancreatitis was induced by 8 hourly intraperitoneal (i.p) injections of caerulein (100 μg/kg body weight) into indicated mice for 2 consecutive days. The mice were then allowed to recover for 7 days or 18 days prior to harvesting pancreatic tissue.

### Western Blotting

Tissues and cells were harvested and lysed with RIPA buffer supplemented with protease and phosphatase inhibitors (MCE). The protein concentration was measured with the BCA Protein Assay (BioRad). The protein was separated by 8%–12% SDS-PAGE gels and transferred onto polyvinylidene fluoride membranes (Millipore). Membranes were blocked in 5% BSA in TBS for 1 hour at room temperature and then incubated with primary antibodies overnight at 4C, washed in TBS containing 0.1% Tween20, incubated with horseradish peroxidase (HRP)-conjugated secondary antibody for 1 hour at room temperature, and developed by ECL reagent (Thermo Fisher Scientific). The primary antibodies used in this study were as follows: NSD2 (Abcam, ab75359), H3K36me1 (Abcam, ab9048), H3K36me2 (Abcam, ab9049), H3K36me3 (Abcam, ab9050), H3 (Cell Signaling Technology, 9715), β-Tubulin (Cell Signaling Technology, 2146), p65 (Cell Signaling Technology, 8242), p-p65(Ser536) (Cell Signaling Technology, 3033), IκBα (Cell Signaling Technology, 4814), p-IκBα(Ser32/36) (Cell Signaling Technology, 9246), IKKα/β (Abcam, ab178870), p-IKKα/β(Ser176/180) (Cell Signaling Technology, 2697).

### Histology and IHC staining

Samples were deparaffinized and rehydrated. Antigen was retrieved using 0.01 M sodium-citrate buffer (pH 6.0) at a sub-boiling temperature for 15 min after boiling in a microwave oven. To block endogenous peroxidase activity, the sections were incubated with 3% hydrogen peroxide for 20 min. After 1 h of pre-incubation in 5% normal goat serum to prevent nonspecific staining, the samples were incubated with the primary antibody against NSD2 (Abcam, ab75359), H3K36me2 (Abcam, ab9049), Ki67 (Abcam, ab6526), CK19 (Abcam, ab15463), p65 (Cell Signaling Technology, 8242) at 4◦C overnight. After three washes in PBS, sections were incubated with an HRP-conjugated secondary antibody for 1 hour at room temperature. Color was developed using the DAB (diaminobenzidine) Substrate Kit (Gene Tech). Counterstaining was carried out using hematoxylin.

### Luciferase reporter assay

The NF-κB firefly luciferase reporter plasmid (Bayotime, Shanghai, China) and renilla luciferase reporter plasmid were co-transfected into 293T cells using Lipofectamine 3000 (Thermo Fisher Scientific). Firefly and Renilla luciferase activities were measured using the Dual-Glo Luciferase Assay System (Promega). Renilla activity was used to normalize luciferase reporter activity. Assays were performed on cells in three wells for each experiment to obtain an average count and in three independent biological replicates.

### Co-immunoprecipitation assays

Cells were washed with cold PBS and lysed in RIPA lysis buffer (50 mmol/L Tris-HCl, pH 8.0; 150 mmol/L NaCl; 1% NP-40) supplemented with protease and phosphatase inhibitors (Millipore) at 24 h after transfection. Cell lysates were incubated with primary antibodies overnight at 4 °C. Pierce™ Protein A/G Magnetic Beads (Thermo) were then added and the lysates were incubated for another 4 h at 4 °C. The immunoprecipitates were washed four times with the lysis buffer and boiled for 5 min at 98 °C in protein loading buffer. Immunoprecipitated proteins were detected by subsequent immunoblotting. Antibodies used in the co-immunoprecipitation experiments were as follows: NSD2 (Abcam, ab75359), p65 (Cell Signaling Technology, 8242), Flag-tag (Cell Signaling Technology, 14793S), HA-tag (Cell Signaling Technology, 3724).

### Chromatin immunoprecipitation sequencing and analyses

Cells were crosslinked with 1 % formaldehyde for 10 min at room temperature and quenched with 125 mM glycine. The fragmented chromatin fragments were pre-cleared and then immunoprecipitated with Protein A + G Magnetic beads coupled with anti-H3K36me2(Abcam, ab9049) or anti-p65 (Cell Signaling Technology, 8242) antibodies. After reverse crosslinking, ChIP and input DNA fragments were end-repaired and A-tailed using the NEBNext End Repair/dA-Tailing Module (E7442, NEB) followed by adaptor ligation with the NEBNext Ultra Ligation Module (E7445, NEB). The DNA libraries were amplified for 15 cycles and sequenced using Illumina NextSeq 500 with single-end 1x75 as the sequencing mode. Raw reads were filtered to obtain high-quality clean reads by removing sequencing adapters, short reads (length <35 bp) and low-quality reads using Cutadapt (v1.9.1) and Trimmomatic (v0.35). Then FastQC is used to ensure high reads quality. The clean reads were mapped to the mouse genome (assembly GRCm38) using the Bowtie2 (v2.2.6) software. Peak detection was performed using the MACS (v2.1.1) [9] peak finding algorithm with 0.01 set as the p-value cutoff. Annotation of peak sites to gene features was performed using the ChIPseeker R package.

### RNA-seq and analyses

For Ctrl and Nsd2-KO of KPC1199 cells, a total of 10^7^ cells were harvested for RNA preparation. Total RNA was extracted from the samples by Trizol reagent (Invitrogen) separately. The RNA quality was checked by Agilent 2200 and kept at −80°C. The RNA with RIN (RNA integrity number) > 7.0 is acceptable for cDNA library construction. The cDNA libraries were constructed for each RNA sample using the TruSeq Stranded mRNA Library Prep Kit (Illumina, Inc.) according to the manufacturer’s instructions. Before read mapping, clean reads were obtained from the raw reads by removing the adaptor sequences and low-quality reads. The clean reads were then aligned to mouse genome (mm10) using the Hisat2. HTseq was used to get gene counts and RPKM method was used to determine the gene expression. We applied DESeq2 algorithm to filter the differentially expressed genes, after the significant analysis, P-value and FDR analysis were subjected to the following criteria: i) Fold Change>1.2 or < 0.833; ii) P-value<0.05. Gene ontology (GO) analysis was performed to facilitate elucidating the biological implications of the differentially expressed genes in the experiment. We downloaded the GO annotations from NCBI (http://www.ncbi.nlm.nih.gov/), UniProt (http://www.uniprot.org/) and the Gene Ontology (http://www.geneontology.org/). Fisher’s exact test was applied to identify the significant GO categories (P-value < 0.05). Pathway analysis was used to find out the significant pathway of the differentially expressed genes according to KEGG database. We turn to the Fisher’s exact test to select the significant pathway, and the threshold of significance was defined by P-value< 0.05.

### Acinar cell explants

Murine pancreas acinar cells were isolated as previously described^52^. The viability of acinar cells was examined by crystal violet staining (>95%). Primary acinar cells were seeded on pre-coated plates with Collagen I (5μg / cm^2^, YEASEN) and cultured in Waymouth medium (Gibco) supplemented with 10% FBS, 1% penicillin/streptomycin and 50 ng/ml TGFa. The ADM (acinar-to-ductal metaplasia) was observed and representative images were taken at the indicated time.

### Electrophoretic mobility shift assay (EMSA)

Nuclear extracts were prepared in KPC1199 cells transfected with indicated vector. The protein concentration from nuclear extracts was detected using the BCA Protein Assay Kit (Thermo). The sequence of biotin-labeled NF-κB p65 probes was 5’-AGTTGAGGGGACTTTCCCAGGC-3’. The DNA-binding activity of NF-κB p65 was determined using the Chemiluminescent EMSA Kit (GS008; Beyotime) as instructed in the manufacturer’s protocol.

### Statistical analysis

All experiments were performed using 3–15 mice or at least three independent repeated experiments. Unless otherwise indicated, data presented as the mean ± S.E.M. All statistical analyses were performed with GraphPad 8.0 software. Student’s t-test assuming equal variance was used, and two-way analysis of variance for independent variance. Pearson correlation coefficients were used to evaluate the relationships between NSD2 and gene expressions. χ2 test were used to determine whether there was a significant difference between the expected frequencies and the observed frequencies in one or more categories. *p < 0.05, **p < 0.01, and ***p <0.001, ****p <0.0001.

### Study approval

All animal experiments were approved by the Renji Hospital Animal Care and Use Committee (202201027). The human pancreatic tumor microarray from Ren Ji Hospital was approved with Local Ethics Committee approval and patient consent (KY2022-036-B).

## Data availability

All data are available from the authors upon reasonable request. RNA-Seq raw data have been deposited in the Gene Expression Omnibus (GEO) under accession number GEO: GSE216562, GSE215386. ChIP-Seq raw data have been deposited in the Gene Expression Omnibus (GEO) under accession number GEO: GSE226058.

## Acknowledgments

This work was supported by funds from National Natural Science Foundation of China (82073104 to L.Li., 82022049 to J.Xue, 82073105 to N.Niu), Science and Technology Commission of Shanghai Municipality (19140905500 to L.Li., 20ZR1432900 to N.Niu), Innovative Research Team of High-Level Local Universities in Shanghai (J.Xue.), Shanghai Municipal Education Commission-Gaofeng Clinical Medicine Grant Support (No. 20161312, J.Xue), State Key Laboratory of Oncogenes and Related Genes (KF2113 to N.Niu) and 111 project (no. B21024).

## Author contributions

L.L., J.X. designed experiment and interpreted data; W.F. performed most of the experiments; P.L., H.R, Z.C., W.Z., C.M, C.L., and Y.X. assisted in some experiments; WQ.G. assisted in some discussion; W.F., L.L. and N.N. wrote the manuscript; L.L., J.X. and N.N. provided the overall guidance.

